# Beyond seizure control: identifying deficits in cognitive networks in absence seizure

**DOI:** 10.1101/2025.09.29.679360

**Authors:** Gil Vantomme, Gabrielle Devienne, Jacob M Hull, John R Huguenard

## Abstract

Absence epilepsy is frequently associated with cognitive impairments, yet the direct or indirect involvement of cognitive circuits remains poorly understood, as spike-and-wave discharges are rarely reported in these regions. Here, we investigate the role of a thalamic–prefrontal pathway in a mouse model of absence epilepsy (Scn8a^+/-^). We find that Scn8a^+/-^ mice exhibit deficits in reversal learning, along with impaired recruitment of medial prefrontal cortex (mPFC) neurons by the Reuniens nucleus of the thalamus. This deficit is accompanied by an altered excitation–inhibition balance and reduced excitability of layer I interneurons in the mPFC, which constitutes the main recipient zone of reuniens inputs to the mPFC. Remarkably, stimulation of Reuniens at 20 Hz significantly reduces seizure incidence and improves performance in reversal learning. Our findings reveal previously unrecognized cognitive circuit dysfunctions in absence epilepsy and highlight the thalamo–prefrontal axis as a promising target for both cognitive and seizure-related interventions.

## Introduction

Typical absence seizures are characterized by sudden, brief lapses of consciousness accompanied by behavioral arrest and stereotypical spike-and-wave discharges (SWDs) (Gibbs et al. 1935; Panayiotopoulos 1999). Beyond the immediate impact of seizures, affected individuals frequently experience psychiatric comorbidities, including deficits in attention, memory, and cognitive flexibility, as well as mood impairments, for review: (Fonseca Wald et al. 2019; Gruenbaum et al. 2021). Importantly, a substantial fraction of patients remains pharmaco-resistant, and comorbidities often persist even when seizures are controlled (D’Agati et al. 2012; Cerminara et al. 2013; Román-Guzmán et al. 2025). These challenges highlight the need to elucidate the mechanisms underlying both seizure activity and associated cognitive deficits, with the goal of developing treatments that address the full spectrum of clinical burden.

The thalamocortical network is a critical driver of typical absence seizures (Steriade et al. 1993; Huguenard and McCormick 2007; Lindquist et al. 2023), yet its impact is not uniformly distributed across brain regions. SWDs generalize rapidly across hemispheres and cortices, with an initial foci in somatosensory and frontal/motor areas reported across species (Crunelli and Leresche 2002; Meeren et al. 2002; Ding and Gallagher 2016; Lee et al. 2019). Limbic structures such as hippocampus show transient and focal increases in BOLD signals during SWD (Nersesyan et al. 2004), increase cerebral glucose utilization rates (Nehlig et al. 1991), and increase synchronicity without full paroxysmal discharge (Velazquez et al. 2007) in typical absence epilepsy, suggesting that they are relatively spared compared to sensory motor areas. This is in contrast with atypical absence epilepsy, where midline thalamus and hippocampus are highly involved in seizure generation for review: (Onat et al. 2013).

This raises the possibility that circuit-specific dysfunctions outside the canonical seizure network contribute to the cognitive comorbidities in typical absence epilepsy. Here, we focused on the Reuniens nucleus of the thalamus (Re). This midline thalamic nucleus forms reciprocal connections with the hippocampus and the mPFC, and is strongly implicated in higher-order cognitive processes, for review: (Dolleman-van Der Weel et al. 2019). Using a combination of *in vitro* and *in vivo* electrophysiological recordings with optogenetic manipulation in the Scn8a^+/-^ mouse model of absence epilepsy, we investigated circuit abnormalities between the Re and mPFC, their impact on SWDs, and cognitive flexibility.

We show that Scn8a^+/-^ mice exhibit impaired cognitive flexibility, recapitulating a key comorbidity observed in patients. Despite minimal engagement of mPFC units in SWDs and the absence of seizure induction by Re stimulation, synaptic recruitment of the mPFC by Re is disrupted. Our data suggest that this deficit arises from reduced feedforward inhibition, in part due to hypoexcitability of mPFC layer 1 interneurons. Strikingly, optogenetic stimulation of Re at 20 Hz reduced seizure incidence *in vivo* and restore performance during reversal learning.

Together, these findings reveal that the Re–mPFC circuit is functionally altered in absence epilepsy and may underlie cognitive comorbidities. They further demonstrate that targeted stimulation of Re is both safe and effective in suppressing seizures, identifying this pathway as a promising candidate for circuit-based therapeutic interventions in patients with refractory epilepsy.

## Results

### SWDs are largely absent in the mPFC of Scn8a^+/-^ mice

The mPFC has been reported as a region involved in SWD during typical absence seizures, either as a propagation site (Lörincz et al. 2007) or even as a focal onset zone in a GHB mouse model (Lee et al. 2019). To examine this in Scn8a^+/-^ mice, we analyzed acute Neuropixels recording from single units in motor cortex (MC), mPFC and olfactory bulbs (OLF) during seizures (Fig. 1A,B; n = 4 mice). We then plotted unit firing rates relative to the phase of the SWDs (Fig. 1C-F). Units in MC exhibited strong phase-locking to the spike component, with a mean vector strength of 0.59 ± 0.01. By contrast, units in mPFC and OLF showed little phase-locking, with mean vector strengths of 0.19 ± 0.02 and 0.12 ± 0.02, respectively (one-way repeated-measures ANOVA: *F*(2,6) = 207.7, *p* = 2.9 x 10^-6^; post hoc paired Student’s *t*-tests with Hochberg correction: MC–mPFC *p* = 0.0003, MC–OLF *p* = 0.0005, mPFC–OLF *p* = 0.11) (Fig. 1G). This lack of phase locking of mPFC units indicates that SWDs are not effectively recruiting or synchronizing neuronal ensembles in this region during ictal events in Scn8a^+/-^ mice, contrasting with regions such as MC, where a strong phase coupling is observed.

**Fig. 1.**
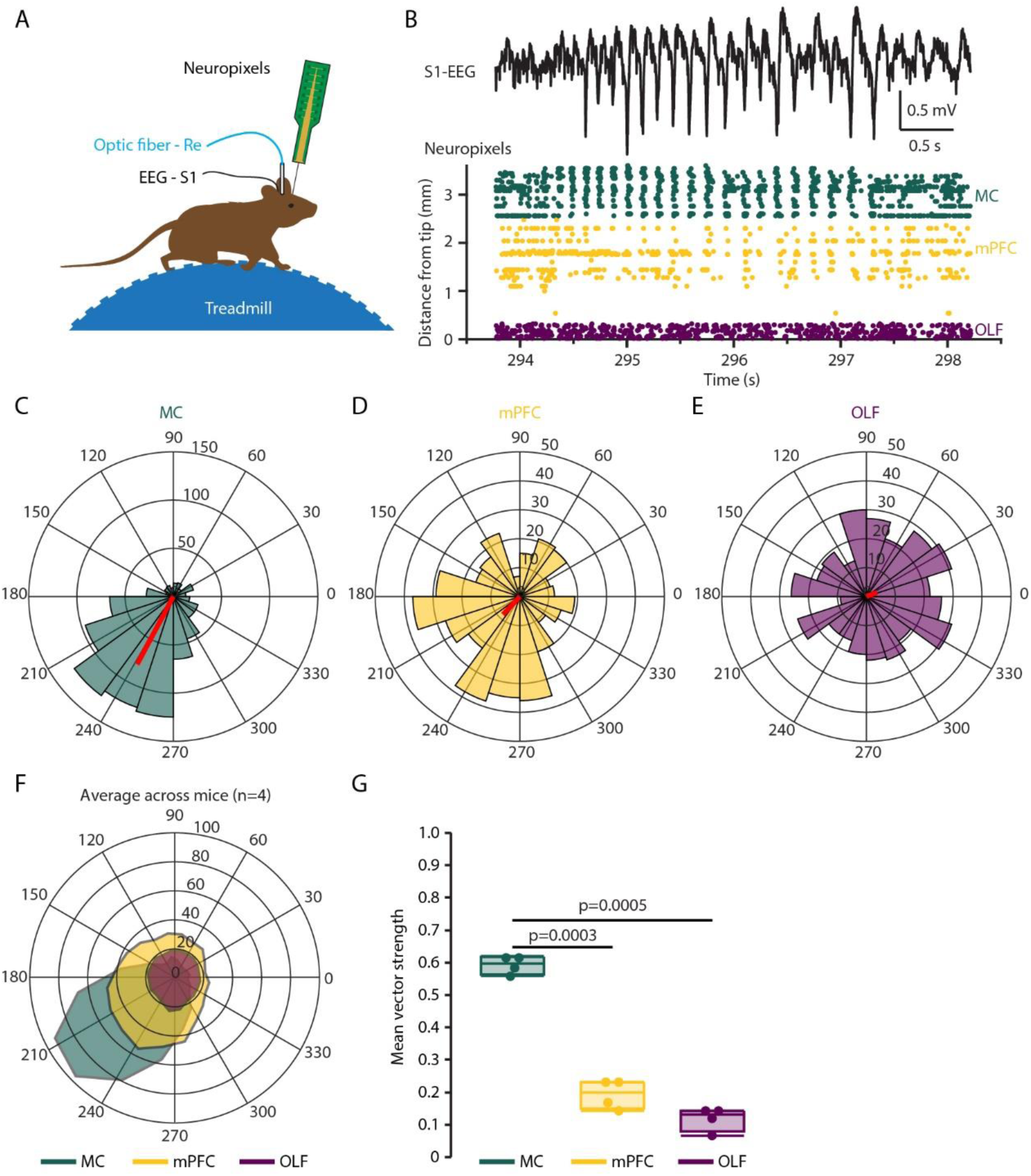
SWDs are largely absent in the mPFC of Scn8a^+/-^ mice. (A) Schematic of the recording setup: acute Neuropixels probe in frontal cortex, EEG in primary somatosensory cortex (S1), and optic fiber targeting the Re. Mice were awake and head-restrained on a cylindrical treadmill. (B) Example recording from a prototypical Scn8a**^+/-^** mouse. SWD is recorded with EEG lead (top) along with corresponding unit activity from the Neuropixels probe in motor cortex (MC), medial prefrontal cortex (mPFC), and olfactory bulb (OLF) (bottom). (C-E) Non-normalized phase plots of unit firing during the recording shown in B for MC (C), mPFC (D), and OLF (E). The red line represents the mean phase angle and non-normalized vector length (F) Average phase plots of unit firing across four mice for MC (green), mPFC (yellow), and OLF (purple). (G) Mean vector strength for analysis in F.

### Re recruitment of the mPFC is altered in the Scn8a^+/-^ mouse model of absence seizures

Although units in the mPFC show little phase-locking to the SWDs, the presence of cognitive deficits in absence epilepsy strongly suggests dysfunction in associated circuits (Ferguson et al. 2023; Fonseca Wald et al. 2019). To test whether the Re–mPFC pathway relevant for cognition is affected in Scn8a^+/-^ mice, we recorded local field potentials (LFPs) in mPFC using a 16-shank linear electrode array spanning cortical layers (Fig. 2A). Slices were obtained from mice expressing ChR2 in Re axons after AAV1-CaMKIIα-ChR2-eYFP injection into Re. Blue light activation of Re afferents evoked LFPs across mPFC layers, from which we derived current source density (CSD) maps (Freeman and Nicholson 1975; Ferguson et al. 2023) (Fig. 2B). In the CSD signals, pairs of sinks and sources were observed following optogenetic stimulation of Re afferents, attributed to cations flowing from the extracellular space into the cells during synaptic excitation, and flowing back out of cells at a distance, respectively. Sources and sinks were prominent within the first 10 ms after the light onset in L2/3 and L5 (Fig. 2C). L2/3 peak CSD sinks (Scn8a^+/+^: -10±2 µA/mm^3^, n=10; Scn8a^+/-^: -10±4 µA/mm^3^, n=8; *F*(1,16) = 0.001, *p* = 0.98) and sources (Scn8a^+/+^: 12±3 µA/mm^3^, n=10; Scn8a^+/-^: -16±5 µA/mm^3^, n=8; *F*(1,16) = 0.43, *p* = 0.52) did not differ between genotypes. By contrast, Scna8^+/-^ showed a significantly larger sink in L5 (Scn8a^+/+^: -3±1 µA/mm^3^, n=10; Scn8a^+/-^: -9±3 µA/mm^3^, n=8; *F*(1,16) = 6.31, *p* = 0.02), indicating altered recruitment of the deeper, output layer of mPFC.

**Fig. 2.**
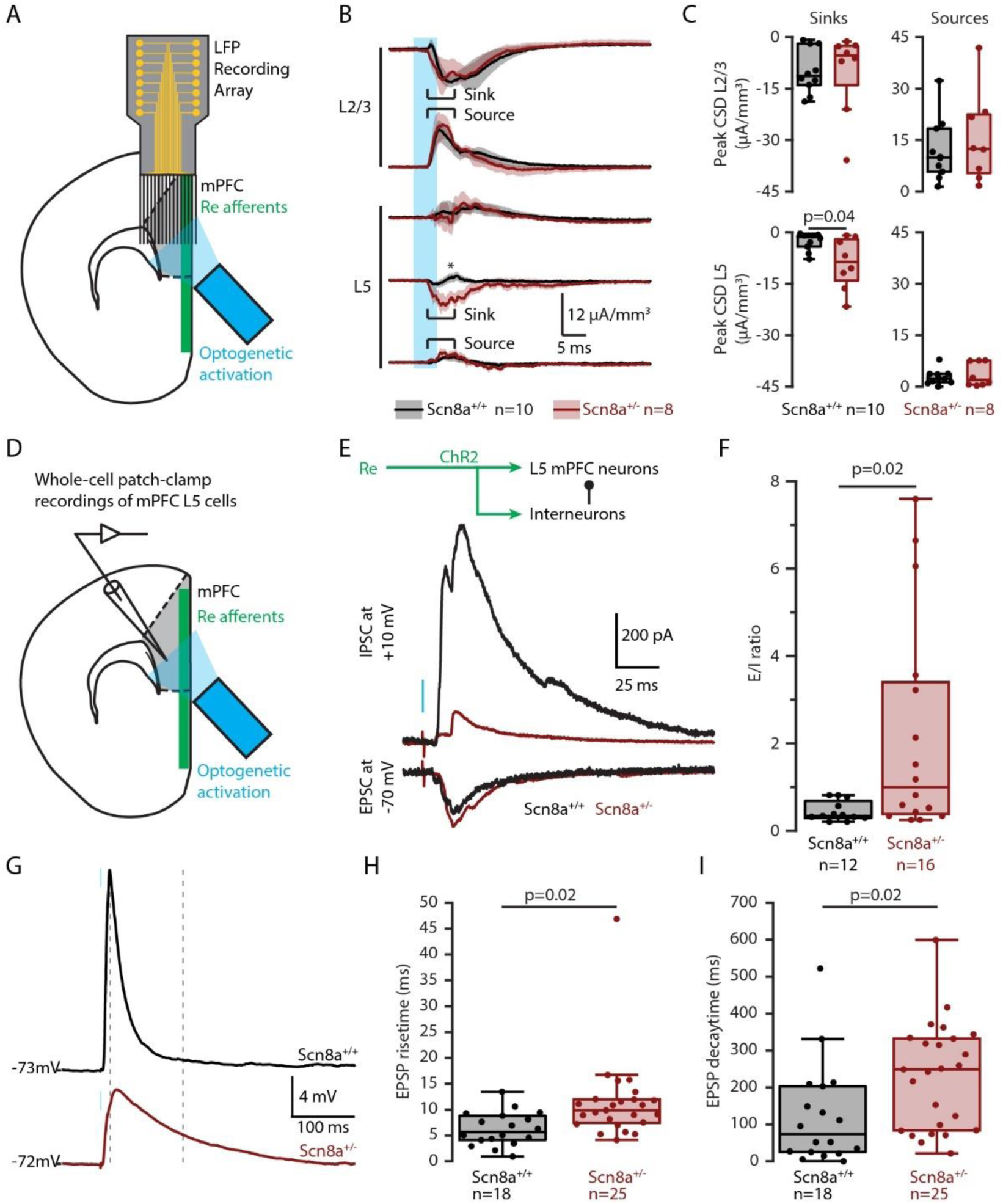
Re recruitment of the mPFC is altered in the Scn8a^+/-^ mice. (A) Recording configuration for local field potentials (LFPs): a 16-channel silicon probe was positioned across cortical layers of the mPFC in acute coronal slices from Scn8a^+/−^ and wild-type littermates expressing ChR2 in Re. Green represents the region of YFP/ChR2 fibers from Re projecting to mPFC L1. (B) Example current source density (CSD) profiles evoked by optogenetic Re activation. (C) Peak CSD sinks and sources in L2/3 (top) and L5 (bottom) of the mPFC. (D) Recording configuration for whole-cell patch clamp: L5 pyramidal neurons in mPFC were recorded in acute slices with ChR2-expressing Re afferents. (E) Example traces of monosynaptic EPSCs (−70 mV) and polysynaptic IPSCs (+10 mV) evoked by Re optogenetic stimulation. (F) Excitatory/Inhibitory (E/I) ratio calculated as EPSC amplitude (−70 mV) divided by IPSC amplitude (+10 mV). (G) Example current-clamp traces showing subthreshold EPSPs evoked by Re stimulation. (H-I) Quantification of EPSP rise time (H) and decay time (I).

To determine the specific circuit alterations underlying the global synaptic source/sink differences, we performed whole-cell patch-clamp recordings from the major output neurons of the mPFC, L5 pyramidal neurons (Fig. 2D). Optogenetic stimulation of Re axons evoked excitatory postsynaptic currents (EPSCs) in L5 cells at –70 mV (near the Cl⁻ reversal potential) and inhibitory postsynaptic currents (IPSCs) at +10 mV (near the AMPAR/NMDAR reversal potential) (Fig 1E). EPSC latencies were short (Scn8a^+/+^: 4.0±0.3 ms, n=12; Scn8a^+/-^: 4.7±0.5 ms, n=16; *F*(1,26) = 1.48, *p* = 0.2), consistent with monosynaptic input, whereas IPSC latencies were longer (Scn8a^+/+^: 8.0±0.5 ms, n=12; Scn8a^+/-^: 10.2±0.9 ms, n=16; *F*(1,26) = 3.92, *p* = 0.06), consistent with polysynaptic recruitment (Suppl. Fig. 1).

Peak EPSC amplitudes were similar across genotypes (Scn8a^+/+^: -331±80 pA; Scn8a^+/-^: -331±68 pA; *F*(1,26) = 0.000, *p* = 1.0). However, IPSC amplitudes were reduced in Scn8a^+/-^ (782±137 pA) compared to their wild-type littermates (422±102 pA; *F*(1,26) = 4.64, *p* = 0.041). This resulted in a marked shift in the excitatory/inhibitory (E/I) ratio: Scn8a^+/+^ mice showed the classical strong feedforward inhibition (E/I = 0.46±0.07), whereas Scn8a^+/-^ cells showed a significantly elevated E/I ratio (2.22 ± 0.62; *F*(1,26) = 5.95, *p* = 0.02), with about half of Scn8a^+/-^ neurons showing little to no feedforward inhibition (Fig. 2F). This synaptic imbalance toward excitation was reflected in prolonged excitatory postsynaptic potentials (EPSPs) recorded in current-clamp at resting membrane potential (Fig. 2G-I). Repetitive 10 Hz stimulation of Re afferents produced short-term depression of both EPSCs and IPSCs in both genotypes (Suppl. Fig. 2).

Because absolute amplitudes of optogenetically evoked currents can vary with ChR2 expression levels, we prepared mPFC slices in which layer 1 (L1), where Re axon are most heavily concentrated (Vertes et al. 2015), was partially disconnected by a cut (Cauller and Connors 1994). A bipolar electrical stimulation electrode was positioned in the semi-isolated L1 flap, while recordings were made from L5 pyramidal neurons in an adjacent intact column (Fig. 3A). This preparation enables precise control of stimulation intensity while assessing how effectively L1 inputs recruit local mPFC circuits (Fig. 3B). In this configuration, Scn8a^+/-^ mice showed similar EPSC amplitudes compared to wild-type littermates (Scn8a^+/+^: -147±46 pA, n=6; Scn8a^+/-^: -99±34 pA, n=10, *F*(1,14) = 0.71, *p* = 0.41). However, IPSCs in Scn8a^+/-^ mice plateaued rapidly with increasing stimulation intensity and were significantly smaller than in controls (Scn8a^+/+^: 356±79 pA, n=6; Scn8a^+/-^: 143±35 pA, n=10, *F*(1,14) = 7.94, *p* = 0.014) (Fig. 3C). These results show that electrical stimulation of mPFC L1 evokes similar monosynaptic EPSC amplitudes in L5 pyramidal neurons across genotypes, but smaller polysynaptic IPSC amplitudes in Scn8a^+/-^ mice, consistent with the findings from optogenetic stimulation of Re afferents (Fig. 2 and Suppl. Fig. 1). This suggests an altered recruitment of inhibitory circuits in the mPFC.

**Fig. 3.**
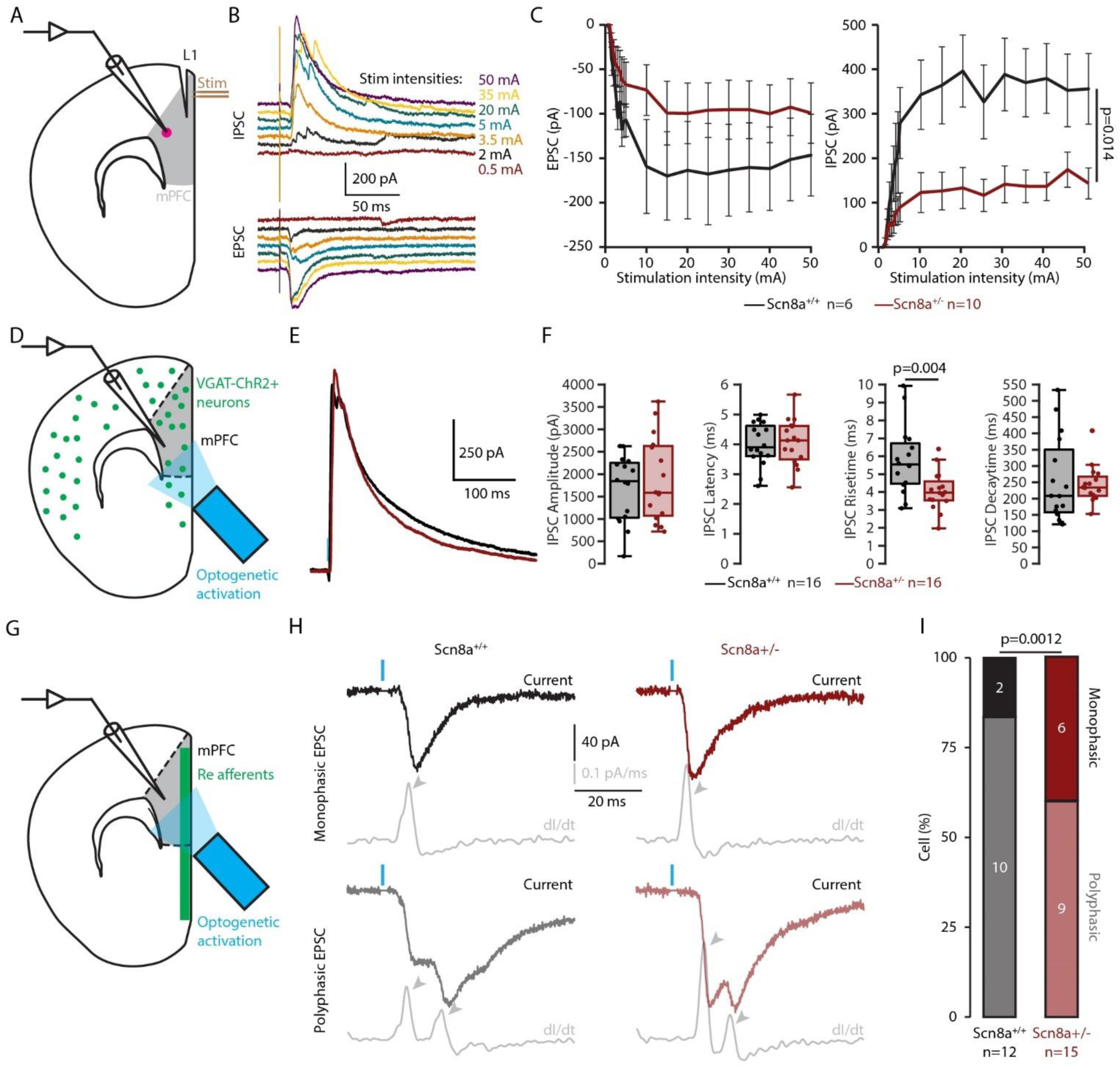
Re recruitment of the mPFC is altered in the Scn8a^+/-^ mice. (A) Whole-cell patch recordings from L5 pyramidal neurons in mPFC slices with a partially disconnected L1 flap. A bipolar electrode delivered electrical stimulation in the L1 flap. (B) Example EPSCs (−70 mV) and IPSCs (+10 mV) evoked by increasing stimulation intensities. (C) Quantification of EPSC and IPSC amplitudes. (D) Whole-cell patch recordings from L5 pyramidal neurons in acute slices from Scn8a^med^-VGAT- ChR2 mice. (E) Example IPSCs evoked by optogenetic activation of local interneurons. (F) Quantification of IPSC properties. (G) Whole-cell patch recordings from L5 pyramidal neurons in slices expressing ChR2 in Re afferents. (H) Example monophasic and polyphasic EPSCs evoked by optogenetic activation of Re afferents. (I) Proportion of L5 neurons responding to Re afferent stimulation with monophasic vs. polyphasic EPSCs.

To test whether this deficit reflected impaired interneuron output, we directly activated local interneurons by optogenetic stimulation in Scn8a^med^-VGAT-ChR2 mice (Fig. 3D). IPSC properties were similar across genotypes (Fig. 3E,F), apart from a slight decrease in rise time in Scn8a^+/-^, suggesting the E/I imbalance in Scn8a^+/-^ arises from reduced polysynaptic recruitment rather than interneurons output deficits.

A particular feature of the Re–mPFC circuit is its propensity to recruit feedforward excitation in addition to the classical feedforward inhibition, as indicated by polyphasic EPSCs (Vantomme et al. 2025). Scn8a^+/-^ mice exhibited a lower proportion of pyramidal cells with polyphasic EPSCs (9/15 cells) compared with wild-type littermates (10/12 cells; χ² test, *p* = 0.001) (Fig. 3G-I). These reductions in both polysynaptic feedforward excitation and inhibition upon stimulation of Re afferents, together with relatively preserved monosynaptic excitation, suggest increased failures in action potential generation or propagation in Scn8a^+/-^ mice, consistent with the NaV1.6 loss-of-function mutation.

### L1 mPFC interneurons are hypoexcitable in Scn8a^+/-^ mice

Beyond the synaptic deficits we observed above, changes in cellular excitability may also contribute to the L5 deficits in the mPFC of Scn8a^+/-^ mice. Previous work showed that activity of mPFC parvalbumin interneurons was affected and directly linked to attentional deficits (Ferguson et al. 2023). Interestingly, L5 pyramidal cells themselves showed little to no changes in excitability (Suppl. Fig. 3). The L1 of the cortex contains a sparse but distinct population of GABAergic interneurons situated at the cortical surface (Chu et al. 2003). These interneurons regulate integration of top-down signals from cortical and subcortical areas, making them critical gatekeepers of input and modulators of deeper-layer activity (Shen et al. 2025). Their ability to filter incoming signals positions them as potential first responders in containing abnormal activity that may precede or trigger seizures.

Whole-cell recordings from L1 interneurons (Fig. 4A) revealed a late-firing pattern in most cases, consistent with prior reports on neurogliaform cells (Shen et al. 2025) (Fig. 4B). Scn8a^+/-^ interneurons showed a higher action potential threshold at rheobase (-35.5±0.9 mV, n=16) compared to wild-type littermates (-38.4±0.9 mV, n=17; *F*(1,31) = 5.21, *p* = 0.03), and a lower action potential afterhyperpolarization (AHP; Scn8a^+/-^: -17.4±0.7 mV, n=16; Scn8a^+/+^: -13.9±1.0 mV, n=17; *F*(1,31) = 7.15, *p* = 0.01) (Fig. 4C,D).

**Fig. 4.**
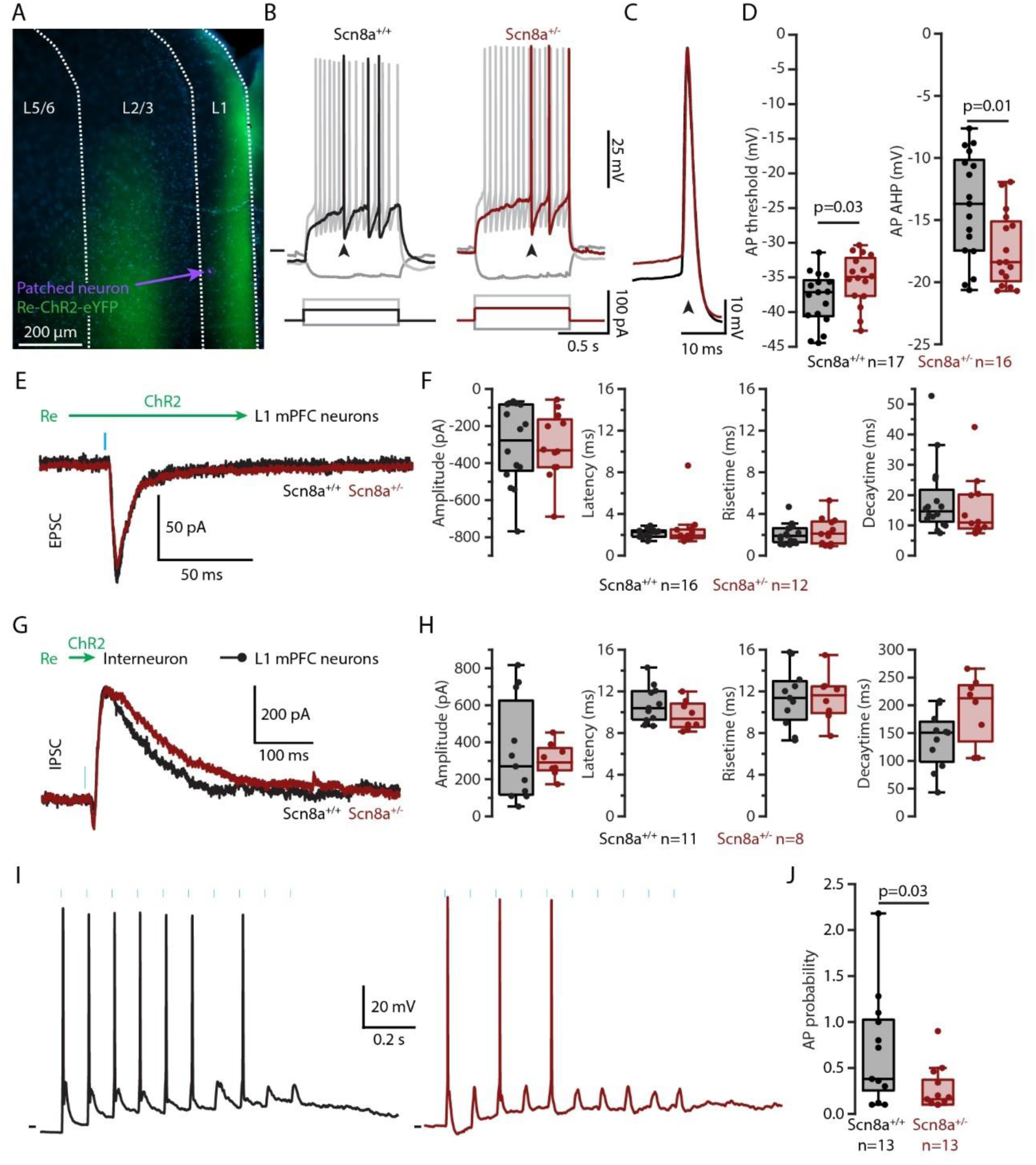
L1 mPFC interneurons are hypoexcitable in Scn8a^+/-^ mice. (A) Epifluorescent micrograph of a mPFC slice showing recorded L1 interneuron (purple), Re afferents expressing ChR2-eYFP (green), and DAPI-stained nuclei (blue). (B) Firing patterns of L1 interneurons. (C) Representative action potentials. (D) Quantification of action potential threshold and afterhyperpolarization. (E) EPSCs recorded in L1 interneurons. (F) Quantification of EPSC properties. (G) IPSCs recorded in L1 interneurons. (H) Quantification of IPSC properties. (I) Action potential discharge from resting membrane potential during 10 Hz train stimulation of Re afferents. (J) Quantification of action potential discharge probability during each stimulus within a 10 Hz train of stimuli.

Optogenetic activation of Re afferents evoked monosynaptic EPSCs in L1 interneurons with comparable latency and kinetics across genotypes (Fig. 4E,F). Feedforward polysynaptic IPSCs were also broadly similar, though Scn8a^+/-^ mice showed a modest increase in decay time (Scn8a^+/-^: 192±22 ms, n=8; Scn8a^+/+^: 138±16 ms, n=11; *F*(1,17) = 4.45, *p* = 0.05) (Fig. 4G,H).

When Re afferents were repeatedly stimulated at 10 Hz, L1 interneurons fired action potentials and subthreshold EPSPs from their resting membrane potential (Fig. 4I). We quantified the number of spikes elicited during a train of 10 stimuli at 10 Hz, considering only recordings in which the first stimulus triggered an AP. Consistent with their elevated threshold, Scn8a^+/-^ interneurons fired significantly fewer action potentials than controls (Scn8a^+/-^: 0.26±0.07, n=13; Scn8a^+/+^: 0.68±0.17, n=13; *F*(1,24) = 5.38, *p* = 0.03) (Fig. 4J).

Together, these findings demonstrate that L1 interneurons in Scn8a^+/-^ mice are hypoexcitable, reducing their ability to respond to Re input and thereby weakening inhibitory control at the cortical entry point.

### Re stimulation reduces seizure incidence in Scn8a^+/-^ mice

Even though there seems to be little to no SWD in limbic structures in typical absence epilepsy, there is evidence of their involvement in seizure modulation (Onat et al. 2013; Dong et al. 2024). We obtained extracellular unit recordings with high density multielectrode silicon probes (Neuropixels). Using such recordings, we confirmed that optogenetic activation of Re outputs *in vivo* induces firing of single units in the mPFC of Scn8a^+/-^ and wildtype littermates (Suppl. Fig. 4). Single spike firing and burst firing were observed in response to Re activation in similar proportions in the mPFC of Scn8a^+/-^ and wildtype littermates (χ² test, p=0.87).

Mice implanted chronically for EEG recordings were trained to rest while head-fixed on a cylindrical treadmill (Fig. 5A). Optogenetic stimulation was 1 min of 20 Hz train followed by 5 min without stimulation, alternating for 1 h (Fig. 5B). The blue light was delivered to the Re through the implanted optic fiber (Re stimulation condition) or delivered inside the chamber as a control (control condition). The number of seizures were counted and averaged across 3 sessions of Re stimulation and 3 sessions of control condition (Fig. 5C-E). Optogenetic activation of Re caused a strong reduction in the seizure incidence compared to control condition over the full 1 h of recording (Fig. 5E; Control: 114±7 seizures/h; Re stim: 67±9 seizures/h; Paired Student’s *t* test, n= 10, *p* = 0.0007) as well as during 1 min-long stimulation blocks (Fig. 5F; Control: 16.8±1.3 seizures/10min; Re stim: 1.5±0.8 seizures/10min; Wilcoxon signed-rank test, n = 10*, p* = 0.006). Altogether, these data show that Re stimulation can negatively modulate seizure incidence in Scna8^+/-^ mice.

**Fig. 5.**
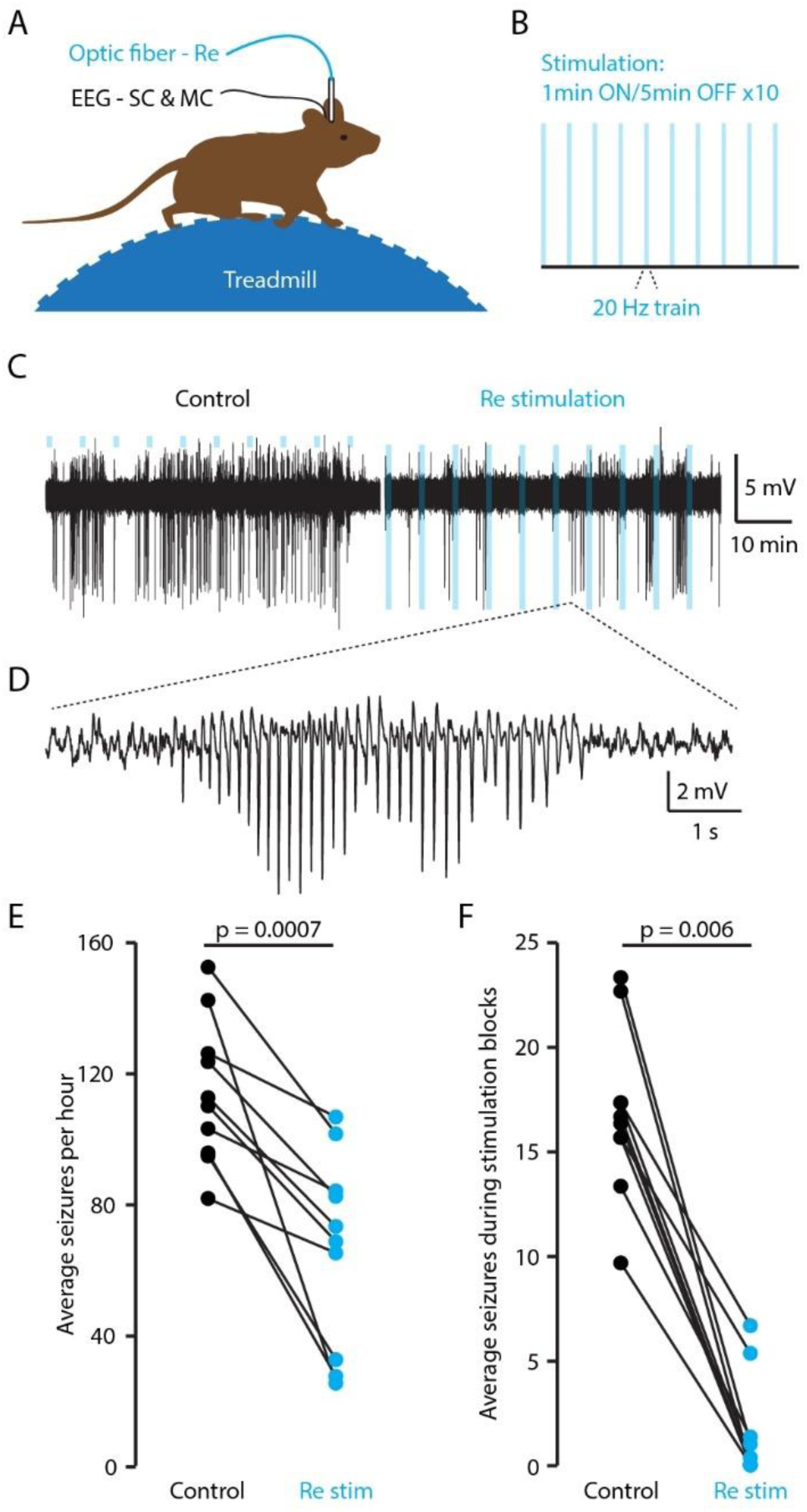
Re stimulation reduces seizure incidence in Scn8a^+/-^ mice. (A) Recording configuration: EEG electrodes in somatosensory cortex (SC) and motor cortex (MC), with optic fiber targeting the Re. Mice were awake and head-restrained on a cylindrical treadmill. (B) Stimulation protocol: ten repetitions of a 1-min 20 Hz train followed by 5 min without stimulation (total duration 1 h). (C) EEG traces from a 1 h recording. Blue bars mark periods of light delivery in the recording cage (left, control stimulation) or on the Re (right). (D) Expanded EEG traces from (C) showing a typical SWD. (E) Average number of seizures during 1 h-long recordings in control and Re stimulation conditions. (F) Same as (E) restricted to the 10 blocks of stimulation.

It has previously been shown that optogenetic excitation or inhibition of thalamic neurons can induce SWD-related seizures (Sorokin et al. 2016; Morais et al. 2025), especially in VPM. By contrast, optogenetic stimulation of Re at 7.5 Hz, the main frequency for absence seizure in Scn8a^+/-^ mice (Papale et al. 2009), did not result in SWDs, consistent with the idea that limbic circuits have little to no involvement in SWD generation (Suppl. Fig. 5).

### Scn8a^+/-^ mice show deficits in cognitive flexibility

Previous work showed that Scn8a^+/-^ mice display impairments in an attentional engagement task (Ferguson et al. 2023). We used a variant of the T-insert within a Morris water maze (Bailoo et al. 2024) to assess cognitive performance in Scn8a^+/−^epileptic mice. This approach allows to measure robust indicators of learning, memory and executive function. It requires a much shorter training period than classical water maze and does not rely heavily on exploration, motivation for novelty or food/water deprivation. Using a Y maze with 120-degree angle avoids sharp turns and makes the maze more natural for mice, reducing learning time (Wijnen et al. 2024). Thus, the swimming Y-maze variant provides a robust behavioral paradigm to detect subtle impairments in spatial learning and flexibility in Scn8a^+/−^ mice. We trained Scn8a^+/-^ and wild-type littermates in the swimming Y-maze task, where mice learned to escape through a fixed tunnel at the end of the correct arm for two consecutive days (10 trials per day; Fig. 6A, left; adapted from (Deacon 2013)). Mice that achieved at least 7/10 successful trials on the second day (escape latency <60s without entering the wrong arm) were advanced to the reversal phase. On the third day, the escape location was switched to the opposite arm (Fig. 6A, right). Approximately half of the mice, regardless of genotype, successfully reach the learning criterion (Fig. 6B). Mice that floated passively during multiple trials without searching for the exit were excluded from analysis (2 Scn8a^+/+^ and 4 Scn8a^+/-^ mice). Scn8a^+/-^ mice and their wild-type littermates performed similarly during initial acquisition of the task (Fig. 6C; escape latency: Scn8a^+/+^: 11.4±2.3 s; Scn8a^+/-^: 14.5±2.4; Student’s *t* test, *p* = 0.75). However, during reversal learning, Scn8a^+/-^ mice displayed significantly longer escape latencies (Scn8a^+/+^: 10.2±1.3 s; Scn8a^+/-^: 23.6±4.3; Dunn test, *p* = 0.008) compared to wild-type littermates (Fig. 6D).

**Fig. 6.**
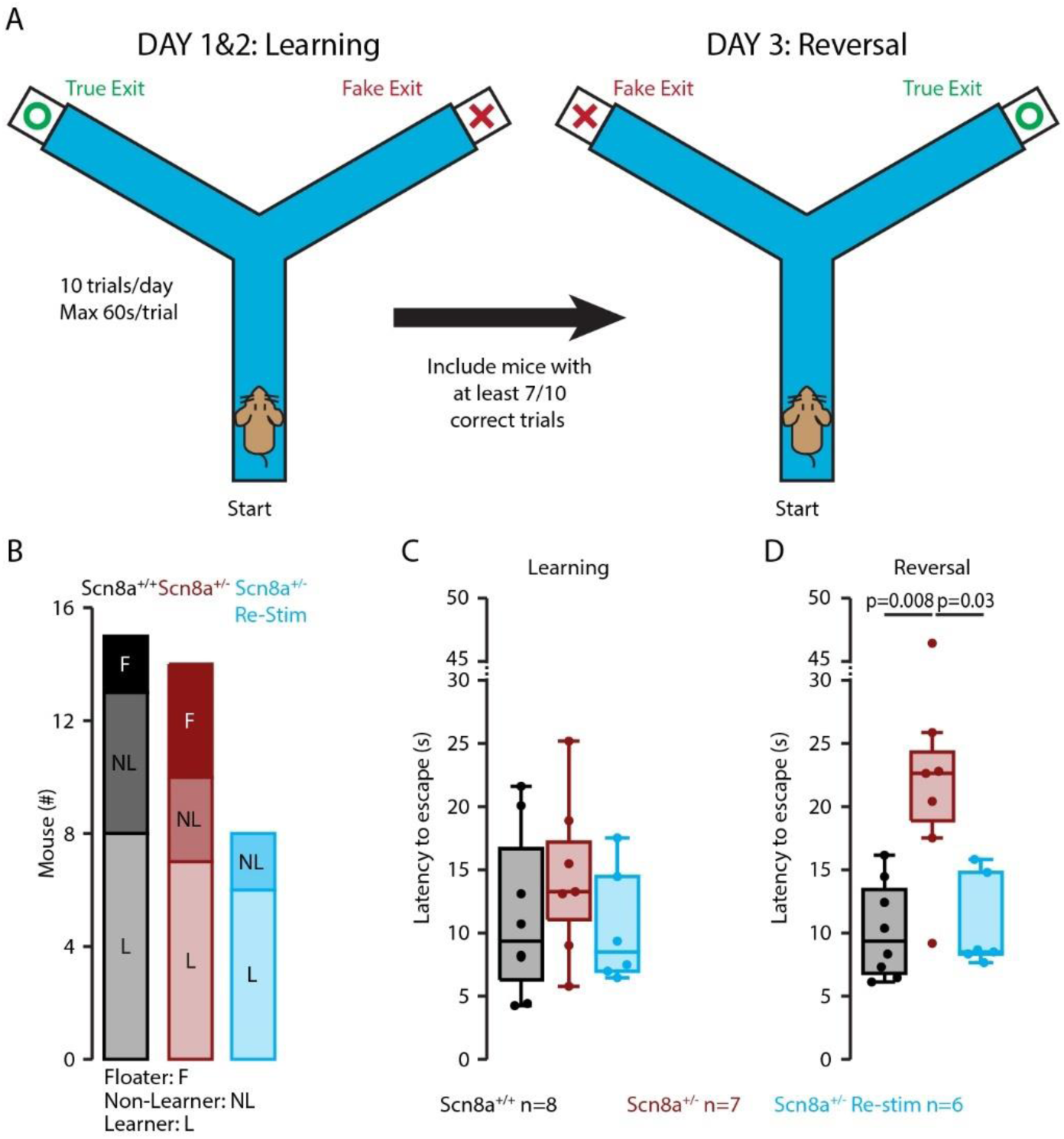
Scn8a^+/-^ mice display perseverant behavior during reversal learning in the swimming Y-maze. (A) Schematic of the swimming Y-maze task. Mice underwent a learning phase on days 1–2 (left) followed by a reversal phase on day 3 (right). (B) Learner (L) mice reached the criterion of ≥7/10 correct trials on day 2 and advanced to the reversal phase. Non- learners (NL) and floaters (F) were excluded from subsequent analyses. (C) Escape latency during the learning phase for Scn8a^+/+^ (black), Scn8a^+/-^ (red) and Scn8a^+/-^ with Re optogenetic stimulation (day 2). (D) Escape latency during the reversal phase (day 3).

To test whether enhancing Re activity could rescue this deficit, Scn8a^+/-^ mice expressing ChR2 in Re received a 20 Hz stimulation train for 1 min each day prior to the 10 trials of the swimming Y maze, as described in Fig. 5. This timing was chosen based on the observation that seizures took some time to resume after the 1 min-long stimulation train, suggesting lasting effect in the thalamocortical networks. These optogenetically stimulated mice (Scn8a^+/-^ Re-Stim) performed similarly to both non-stimulated Scn8a^+/-^ and wild-type littermates during the task acquisition (escape latency: 10.4±1.8 s; one-way ANOVA: *F*(2,18) = 0.84, *p* = 0.45; Fig. 6C). Interestingly, during reversal learning, optogenetically stimulated Scn8a^+/-^ mice showed escape latencies (10.7±1.5 s) comparable to wild-type littermates, thereby rescuing the performance deficit observed in non-stimulated Scn8a^+/-^ mice (Kruskal-Wallis test *p*= 0.0055; post hoc Dunn tests with Hochberg correction: Scna8^+/+^ vs. Scn8a^+/-^ *p* = 0.008; Scna8^+/+^ vs. Scn8a^+/-^ Re-Stim *p* = 0.73; Scn8a^+/-^ Re-Stim vs. Scn8a^+/-^ *p* = 0.026; Fig. 6D).

Together, these results demonstrate that Scn8a^+/-^ mice retain intact learning capacity but exhibit impaired cognitive flexibility, and that optogenetic stimulation of Re prior to testing rescues reversal learning performance to wild-type levels without affecting acquisition.

## Discussion

Uncontrolled seizures in childhood carry devastating long-term consequences, including cognitive decline, psychiatric comorbidities, psychosocial disability, and premature mortality (Sperling 2004). Because of the strong bidirectional links between epilepsy, cognition, and psychiatric health, interventions that reduce seizures may simultaneously improve broader neurodevelopmental outcomes. Thalamic neuromodulation has already demonstrated efficacy for drug-resistant epilepsy (Fisher 2023), yet there is no consensus on the optimal stimulation target for generalized seizures, leaving a critical unmet need. The Re nucleus of the thalamus offers a compelling new avenue: strategically positioned at the interface of the hippocampus and mPFC, it is integral to cognitive control and memory processes (Dolleman-van Der Weel et al. 2019). Emerging evidence, including our findings and those of others (Dong et al. 2024), suggests that modulating Re activity may not only suppress seizures but also address the cognitive comorbidities that burden patients with absence epilepsy.

Previous work demonstrated how defects in mPFC PV interneurons contributed to attention deficits in this mouse model (Ferguson et al. 2023). We expand here by showing that the Scn8a^+/-^ mice also show deficits in cognitive flexibility (Fig. 6), alteration of Re recruitment of mPFC with excitation/inhibition imbalance (Fig. 2 and 3) and hypoexcitability of L1 interneurons (Fig. 4). We also show that optogenetic stimulation of Re does not trigger seizures even when applied at the seizure basic frequency (Suppl. Fig 5), which contrasts with equivalent stimulation in sensory thalamus (Sorokin et al. 2016; Morais et al. 2025). By contrast, Re stimulation at 20 Hz reduces seizure incidence and rescues reversal learning performance, highlighting its potential for neuromodulation.

Our findings are limited to a single mouse model of typical absence epilepsy, the Scn8a^+/−^ line, in which a heterozygous loss-of-function mutation in the NaV1.6 channel is thought to drive hypersynchronous activity between the thalamic reticular nucleus and sensorimotor thalamus, propagating to the cortex as spike-and-wave discharges (Makinson et al. 2017). Notably, selective loss of NaV1.6 within the reticular nucleus alone is sufficient to produce absence seizures, likely through a combination of intrinsic and synaptic impairments in this structure. Beyond the direct channelopathy, Scn8a^+/−^ mice also exhibit maladaptive myelination changes secondary to absence seizures, which further promote epilepsy progression (Knowles et al. 2022). Together, these sodium channel and myelination deficits may contribute to the altered excitability and excitation–inhibition imbalance we observed in the Re–mPFC pathway (Fig. 2 and 3). However, it remains important to recognize that other genetic or pharmacological models of absence epilepsy, while converging on abnormal thalamocortical synchrony and spike-and-wave discharges, may not share the same circuit-level alterations seen in Scn8a^+/−^ mice. Extending these investigations across additional models will therefore be essential to distinguish model-specific features from mechanisms that are conserved across absence epilepsies.

Similarly, our study was restricted to the thalamocortical pathway linking the Re and mPFC. Other higher-order thalamic nuclei, including ventromedial (Paz et al. 2007), centromedial (Zillgitt et al. 2022) and anterior thalamic nuclei (Venkatesh et al. 2023), have also been implicated in seizure modulation. Moreover, the Re sends projections to several additional regions, such as the entorhinal cortex and hippocampus reviewed in (Cassel et al. 2021), which have not yet been examined for synaptic or cellular abnormalities in Scn8a^+/−^ mice. Dysfunctions within these circuits may also contribute to the cognitive impairments we observed.

The mechanisms underlying the seizure-reducing effect of Re stimulation remain to be fully defined. A knock-in model of human BK-D434G channelopathy recapitulates absence seizures with behavioral arrests and shows increased early gene activation (c-Fos) and hypersynchronous bursting in midline thalamic nuclei, including Re (Dong et al. 2022; 2024). Suppressing this midline bursting via pharmacological, optogenetic, or electrical interventions reduced seizure generation and enhanced vigilance, paralleling effects observed with physical perturbation or psychostimulants. These data suggest that midline thalamic nuclei may actively contribute to absence seizure initiation and propagation. Notably, seizure incidence is highest during quiet wakefulness and drowsy states, whereas heightened vigilance suppresses seizures (Bureau et al. 1968; Ferguson et al. 2023). Given the established role of the centromedial thalamus in arousal and attention (Ilyas et al. 2019), midline stimulation may act through two converging mechanisms: disrupting pathological synchronization within thalamocortical circuits and enhancing vigilance, both of which would be expected to reduce seizure incidence. Consistent with this, 20 Hz optogenetic stimulation centered on Re decreased seizure frequency in Scn8a^+/-^ mice (Fig. 6). However, we did not observe a uniform increase in locomotor activity across our cohort. This variability may reflect subtle differences in viral targeting or optic fiber placement, with possible co-stimulation of adjacent centromedial and ventromedial thalamus.

The mechanisms underlying the rescue of reversal learning deficits observed after Re stimulation in Scn8a^+/−^ mice remain unclear and warrant further study. Our recordings (Fig. 5) revealed that seizure reduction persisted beyond the stimulation period, indicating that trains of Re activation may exert lasting effects on thalamocortical networks. Such effects could result from a combination of acute neural mechanisms, such as depolarization block (Lowet et al. 2022) or desynchronization of pathological oscillations as seen in Parkison’s disease (Eusebio et al. 2012), and longer-term adaptive changes in synaptic strength (Xu et al. 2023). Given its connectivity with both the mPFC and hippocampus, the Re has been shown to coordinate activity in its postsynaptic targets at beta frequencies (15–30 Hz) during sequence memory retrieval (Jayachandran et al. 2022). It is therefore plausible that stimulation of Re within this frequency range enhances coherence between the mPFC and hippocampus, potentially serving as a neurobiological substrate for an “episodic buffer” that transiently maintains information stability in the period preceding a decision (Baddeley 2000; Jayachandran et al. 2022).

In summary, our findings demonstrate that the Re targeted activation can reduce seizure incidence without exacerbating ictogenesis and rescue performance in reversal learning. By linking alterations in Re–mPFC circuitry, interneuron excitability, and cognitive flexibility deficits in *Scn8a*^+/-^ mice, we extend the role of the Re beyond cognition into seizure modulation and cognitive restoration. These results support the Re as a promising target for neuromodulation strategies in generalized epilepsies, while also emphasizing the need to refine stimulation parameters and assess circuit specificity. Future work should explore whether different stimulation frequencies or approaches to selectively recruit interneuron populations can optimize both seizure control and cognitive outcomes, paving the way toward translational strategies for pediatric drug-resistant epilepsy.

## Materials and Methods

### Experimental model and subject details

All experimental procedures were performed in accordance with the guidelines of the Institutional Animal Care and Use Committees of Stanford University. This study used young adult (8 to 15 weeks old) Scn8a^med^ mice (C3Fe.Cg-Scn8amed/J, Jackson Laboratory, RRID: IMSR_JAX:003798) and Scn8a^med^-VGAT-ChR2+ mice obtained from crossing Scn8a^+/-^ with hemizygous VGAT-ChR2-EYFP mice (B6.Cg-Tg(Slc32a1-COP4*H134R/EYFP)8Gfng/J, Jackson Laboratory, RRID:IMSR_JAX:014548). Mice were housed in a temperature and humidity-controlled animal house, maintained in a 12 h/12 h light-dark cycle, and could access food and water ad libitum.

### Viral injections

Young adult mice (P30 – 60) were anesthetized with isoflurane (5% induction, 2% maintenance) and received Carprofen 10 mg/kg subcutaneous. A volume of 0.5 µl of virus encoding the ChR2 (AAV1-CaMKIIa-ChR2(H134R)_eYFP-WPRE-HGH, 10^12^ GC, 100 nl/min, Penn Vector Core, Addgene 26969P) was delivered in Re (in mm from bregma, AP -0.75, ML 0, DV -4).

### *In vitro* patch-clamp recording

Acute brain slices were prepared from mice following established protocols (Vantomme et al. 2025). Coronal slices (300 µm thickness) encompassing the medial prefrontal cortex (mPFC) were obtained using a vibratome (Leica VT1200S). The targeted region spanned +1.4 to +2.0 mm anterior to bregma, and recordings were performed within the prelimbic and infralimbic subregions.

Slicing was carried out in an ice-cold, oxygenated solution containing (in mM): 66 NaCl, 2.5 KCl, 1.25 NaH₂PO₄, 26 NaHCO₃, 105 sucrose, 27 glucose, 1.7 ascorbic acid, 0.5 CaCl₂, and 7 MgCl₂. Slices were then incubated at 35°C for 30 minutes in a recovery solution, followed by at least 30 minutes at room temperature prior to recording. The recovery solution contained (in mM): 131 NaCl, 2.5 KCl, 1.25 NaH₂PO₄, 26 NaHCO₃, 20 glucose, 1.7 ascorbic acid, 2 CaCl₂, 1.2 MgCl₂, 3 myo-inositol, and 2 pyruvate. During recordings, slices were perfused with artificial cerebrospinal fluid (ACSF) composed of (in mM): 131 NaCl, 2.5 KCl, 1.25 NaH₂PO₄, 26 NaHCO₃, 20 glucose, 1.7 ascorbic acid, 2 CaCl₂, and 1.2 MgCl₂. Two intracellular solutions were used depending on the experimental configuration. The cesium-based solution consisted of (in mM): 127 Cs- gluconate, 10 HEPES, 2 Cs-BAPTA, 6 MgCl₂, 10 phosphocreatine, 2 Mg-ATP, 0.4 Na-GTP, and 2 QX-314-Cl (Tocris, 2313), adjusted to pH 7.3 and 290–305 mOsm. The potassium-based solution contained (in mM): 144 K-gluconate, 10 HEPES, 3 MgCl₂, and 0.5 EGTA, with the same pH and osmolarity.

Series resistance (Rs) and input resistance (Ri) were monitored throughout the experiments using brief voltage or current pulses, depending on the recording mode. Data were excluded if Rs or Ri varied by more than 20% over the course of the recording. No correction was applied for the measured -10 mV liquid junction potential.

Passive membrane properties were assessed in voltage clamp at a holding potential of -70 mV using 500 ms hyperpolarizing voltage steps (10 mV), or alternatively, in current clamp from the resting membrane potential. Action potential characteristics were evaluated at rheobase under current-clamp conditions, using 500 ms depolarizing current injections of increasing amplitude.

Channelrhodopsin-2 (ChR2)-expressing afferents were stimulated using blue light (1 ms duration) delivered to the full microscopic field via either epifluorescent illumination with an LED light source (Thorlabs M450LP2, 450 nm, 13 mW, 19 mW/cm²) or a laser (Laserglow Technologies, 473 nm, 8 mW, 10.8 mW/cm²). To isolate excitatory and inhibitory synaptic currents, mPFC neurons were held in voltage clamp at -70 mV and +10 mV, respectively.

### *In vitro* local field potential recordings

Extracellular field recordings were performed using procedures previously described (Vantomme et al. 2025). Coronal brain slices (400 µm thick) containing the mPFC were prepared from mice expressing ChR2-eYFP in the nucleus reuniens (Re) following viral injection. Slices were placed in a continuously oxygenated, humidified interface recording chamber maintained at 34°C and perfused with oxygenated artificial cerebrospinal fluid (ACSF) at a flow rate of 2 ml/min.

Local field potentials (LFPs) were recorded using a linear 16-channel silicon probe (NeuroNexus Technologies) with 100 µm inter-electrode spacing, oriented perpendicular to the cortical laminae to span the full depth of the mPFC. Signals were amplified and digitized using a PZ5-32 preamplifier (Tucker-Davis Technologies, TDT) and processed with an RZ5D multichannel processor (TDT). Acquisition protocols were implemented using the Real-time Processor Visual Design Studio (RPvdsEx) along with custom-written Python scripts. Data were sampled at 25 kHz and filtered between 0.1 Hz and 500 Hz.

Optogenetic activation of ChR2-expressing axons was achieved using a 473 nm blue laser (Laserglow Technologies; 5 ms pulse duration; max output 15 mW, 20 mW/cm²). The light was delivered in circular spots approximately 1 mm in diameter, targeting all cortical layers of the mPFC. Stimulation consisted of single pulses delivered every 20 seconds.

To dissect the synaptic contributions to the LFP, pharmacological agents were sequentially applied to the bath. GABA_A_ receptor-mediated inhibition was blocked with Gabazine (Abcam, ab120042; 10 µM), AMPA receptors with DNQX (Abcam, ab120169; 40 µM), NMDA receptors with APV (Sigma, A5282; 100 µM), and voltage-gated sodium channels with tetrodotoxin (TTX, Latoxan L8503; 0.25 µM). Each condition (baseline ACSF, Gabazine, DNQX/APV, and TTX) was recorded for 5–8 minutes, and the final 2 minutes of each recording were analyzed to ensure steady-state drug effects.

To localize and quantify current flow associated with network activity, we computed the one-dimensional current-source density (CSD) from the LFP signals (Freeman and Nicholson 1975). Assuming a uniform extracellular conductivity of 0.3 S/m (Goto et al. 2010), CSD was calculated by applying a second-order spatial derivative to signals from adjacent electrodes. The resulting sink and source patterns reflect transmembrane current flow within specific cortical layers. The successive pharmacological blockade of synaptic and action potential-mediated transmission allowed to disentangle the origins of the recorded sinks and sources in mPFC.

### Chronic *in vivo* EEG recordings

Scn8a^+/-^ mice injected with AAV1-CaMKIIa-ChR2 into the Re were surgically implanted with an optical fiber (200 µm core diameter, 0.22 NA, 2.5 mm ceramic ferrule; Thorlabs) positioned 200 µm above the injection site. For seizure monitoring, two gold pins (1.27 mm, Mill-Max Manufacturing Corp.) were placed in contact with the surface of the primary somatosensory and motor cortices, serving as EEG electrodes. A third gold pin was implanted into the occipital bone to function as both ground and reference.

A custom metal head bar (eMachineShop) was placed and stabilized using dental cement (Metabond). Following surgery, mice were given one week to recover, after which they underwent 2–3 weeks of habituation to handling and head restraint on a cylindrical treadmill.

After a 1-h period of quiet resting on the cylindrical treadmill, mice were recorded for 1 h under two conditions: (1) the test condition, during which laser stimulation was delivered to the Reuniens, or (2) the control condition, during which the laser was not connected to the optic fiber implanted on the mouse head. For stimulation, blue laser pulses (473 nm; Laserglow Technologies) were delivered in 1-min trains at 20 Hz (5-ms pulse duration, 50-ms pulse interval, 3.9 mW output, 3.4 mW/cm²), interleaved with 5-min intervals without stimulation, and repeated throughout the recording period. Recordings were acquired using OpenEphys software (https://open-ephys.org/) at a sampling rate of 30 kHz.

Each mouse underwent three sessions in each condition (Re stimulation vs Control). The seizure frequency of each session was quantified, and average seizure rates were compared between the two conditions.

### Acute *in vivo* Neuropixels recordings

#### Mouse preparation

Experiments were conducted on four Scn8a^+/-^ and three Scn8a^+/+^ mice that had received stereotaxic injections of AAV1- CaMKIIa-ChR2 into the Re. An optical fiber (200 µm core diameter, 0.22 numerical aperture, 2.5 mm ceramic ferrule; Thorlabs) was implanted 200 µm above the viral injection site. A stainless steel miniature self-tapping screw (J.I. Morris Company, FF00CE125), connected to a Mill-Max pin (853-93-100-10-001000), was placed into the occipital bone over the cerebellum to serve as a reference electrode. To monitor cortical seizure activity, a 1.27 mm gold pin (Mill-Max Manufacturing Corp.) was positioned in contact with the surface of the primary somatosensory cortex to function as an EEG electrode. A custom-fabricated metal head bar (eMachineShop) was affixed to the skull and stabilized with dental cement (Metabond). Following surgery, mice were allowed to recover for one week and then gradually acclimated to handling and head-fixation while walking on a cylindrical treadmill. Electrophysiological recordings were performed after 2–3 weeks of habituation.

On the recording day, a 1 mm diameter craniotomy was performed over the right motor cortex (AP: +1.5 to +2.5 mm; ML: +1 to +2 mm) using a 0.5 mm burr (Fine Science Tools). The exposed brain surface was protected with a removable silicone elastomer (Kwik-Cast; World Precision Instruments), and animals were returned to their home cages for at least 4 hours before recordings began. During recording sessions, mice were head-fixed on the treadmill, and the pre-implanted optical fiber was connected to a 473 nm laser (OEM Laser Systems; 5 ms pulse duration; maximum output 3.2 mW, 4.5 mW/cm²). EEG signals were recorded via the cortical gold pin and cerebellar reference screw using an RHD2132 headstage (Intan Technologies) and OpenEphys acquisition system. Immediately prior to probe insertion, the silicone covering the craniotomy was removed. A Neuropixels 1.0 probe, pre-coated with a red fluorescent dye (Vybrant DiD Cell Labeling Solution; ThermoFisher Scientific), was mounted on a cable connected to a PXIe acquisition platform (National Instruments, NI-PXIe-1071) and carefully positioned above the cortical surface for recording.

#### Recording

The Neuropixels probe was slowly lowered into the brain at a 45° angle over a ∼5-minute period using a Sutter MP-285 micromanipulator. The insertion targeted a 3.8 mm span encompassing the motor cortex and mPFC, crossing the midline. After insertion, the probe was allowed to stabilize for approximately 20 minutes before beginning data acquisition. Each recording session consisted of a 5-minute baseline period, followed by 5 minutes of optogenetic stimulation at either 1 Hz or 10 Hz. Stimulation trains were delivered every 10-20 seconds and included either single pulse, 2 pulses (at 1 Hz) or 11 pulses (at 10 Hz), each pulse lasting 5 milliseconds. Signals were recorded at a 30 kHz sampling rate using the OpenEphys acquisition system (https://open-ephys.org/).

#### Analysis

Spike sorting was performed automatically using Kilosort2.5 (https://github.com/MouseLand/Kilosort) followed by manual curating using Phy2 (https://github.com/cortex-lab/phy). Single-unit clusters were excluded if they exhibited inter-spike interval (ISI) violations exceeding 0.05%, defined as the proportion of spikes occurring within the ±1.5 ms refractory window relative to the total spike count. Units that showed instability across the full 10-minute recording session were also discarded from further analysis. Peri-stimulus raster plots and peri-event time histograms were generated using custom-written MATLAB scripts (www.mathworks.com). Z-scores were calculated using the pre-stimulation baseline period (−200 to 0 ms) as the reference distribution.

Absence seizures were identified visually based on the characteristic EEG pattern, consisting of brief, sharp spike components followed by slower wave activity. To analyze the relationship between neuronal firing and the rhythmic seizure oscillation, the EEG signal was band-pass filtered between 6 and 8 Hz to isolate the dominant spike-and-wave frequency. The instantaneous phase of the seizure oscillation was extracted by applying a Hilbert transform to the filtered EEG signal, yielding a continuous phase angle time series ranging from 0° to 360°. Each spike recorded from single units during seizures was then assigned a corresponding phase value based on its timing within this phase time series. To quantify the phase preference of neuronal firing, the seizure phase cycle was divided into 20° bins, and spike counts were computed within each bin to generate phase histograms representing firing probability across the oscillatory cycle. For each seizure event, circular statistics were employed to calculate the resultant vector length and mean vector strength. The mean vector strength is a normalized version of the mean vector length reflecting phase synchrony, with values near zero indicating uniform firing across all phases, and values near one indicating strong phase preference. These measures were computed individually for each seizure and then averaged across seizures within each animal to obtain representative phase-locking metrics per mouse.

To assess whether optogenetic stimulation of the Re could trigger absence seizures, we quantified the temporal relationship between each light pulse and subsequent seizure initiation. Specifically, for each stimulation train, we calculated the latency to the closest following seizure by measuring the time interval between the onset of the stimulation and the start of the nearest seizure event. Seizure occurrence was then binned using 0.5-second intervals to generate a histogram of seizure probability as a function of time following stimulation. This approach allowed us to evaluate whether seizures were more likely to occur shortly after stimulation, as would be expected if stimulation of Re promoted seizure generation.

#### Histology and probe localization

At the conclusion of experiments, mice were euthanized via intraperitoneal injection of Fatal+ (pentobarbital; Vortech Pharmaceuticals), followed by transcardial perfusion with 4% paraformaldehyde (PFA) in phosphate-buffered saline (PBS). Brains were post-fixed overnight in 4% PFA, rinsed in PBS, and sectioned at 100 µm thickness using a vibratome (Leica VT1000S). Sections containing the Neuropixels recording site in the mPFC and the viral injection site in the Re were mounted using DAPI-Fluoromount-G (Southern Biotech, 0100-20). Fluorescent images were acquired on a Zeiss Axio Imager M2 microscope to visualize DAPI staining, ChR2-eYFP expression, and the fluorescently labeled probe track. Neuropixels probe localization was reconstructed in three dimensions using SHARP-track (Allen CCF tools), which enables alignment of histological sections and electrophysiological landmarks to the Allen Mouse Brain Common Coordinate Framework. Based on this alignment, each recording channel was assigned to its respective anatomical location within either the motor cortex, mPFC, or olfactory bulb. For anatomical attribution, the mPFC was defined to include the orbital, prelimbic, and infralimbic cortices, the anterior cingulate cortex and frontal pole, corresponding to areas 24, 25, and 32 according to the Paxinos and Franklin Mouse Brain Atlas, 5th edition.

### Swimming Y maze test

The Y maze consisted of three transparent Plexiglas arms, each 28 cm long, 8 cm wide, and 20 cm high, arranged at 120° angles. The maze was filled with water maintained at 20–22°C. Two of the arms were perforated and connected to black plastic pipes, with the water level adjusted to reach the base of the pipes. One pipe was occluded with a transparent barrier (false exit), while the other remained open (true exit). The escape pipe could be detached and used to transfer the mouse to a heating pad between trials.

Distant visual cues were present in the room to facilitate spatial orientation: black-and-white horizontal stripes on the left wall (60 cm from the maze), a black curtain on the right, and a white rear wall.

All mice began from the same starting arm. The escape location was fixed to either the left or right arm during the first two sessions (learning phase). On the third day (reversal phase), the escape location was switched to the opposite arm. Each session included ten trials, in which the mouse was given 60 seconds to find the correct escape route. Between trials, mice were transferred to a warmed recovery cage for 10–15 seconds. Scn8a^+/-^ mice expressing ChR2 in Re were optogenetically stimulated at 20 Hz for 1 min prior to the block of 10 trials, each day.

A trial was considered successful if the mouse escaped without entering the incorrect arm. It was marked as a failure if the mouse entered the wrong arm or failed to escape within 60 seconds. Mice that floated passively during multiple trials without actively searching for the exit were excluded from analysis (2 Scn8a^+/+^ and 4 Scn8a^+/-^ mice). An additional 5 Scn8a^+/+^ and 3 Scn8a^+/-^ mice were excluded for not meeting the learning criterion (≥7 successful trials on Day 2). The primary behavioral metrics analyzed were the average escape latency and number of arm entries.

### Quantification and statistical analysis

All statistical analyses were performed using 2024 JMP® 18.0.2 (JMP Statistical Discovery LLC, Cary, NC) and the R programming software (3.6.1, The R Foundation for Statistical Computing, 2019). Comparisons between groups were assessed using least squares regression models with fixed effects or multivariate analysis of variance (MANOVA). Statistical significance was determined by F-tests on the least squares means. Post hoc comparisons were corrected for multiple testing where appropriate. Data are reported as mean ± SEM, and significance was set at p < 0.05. Significant exact p values are indicated in the main text and on figures.

## Acknowledgments

We are grateful to Prof. Anita Lüthi for generously providing laboratory space, access to equipment, and overall support during the early stages of this project. We also thank all members of the lab for their valuable feedback on the manuscript and for engaging discussions throughout the course of the study. Special thanks to Nicole Agranonik, Cameron Glick, and Katlin Villar for their assistance with mouse line maintenance, equipment management, and laboratory logistics.

---

**Suppl. Fig. 1.**
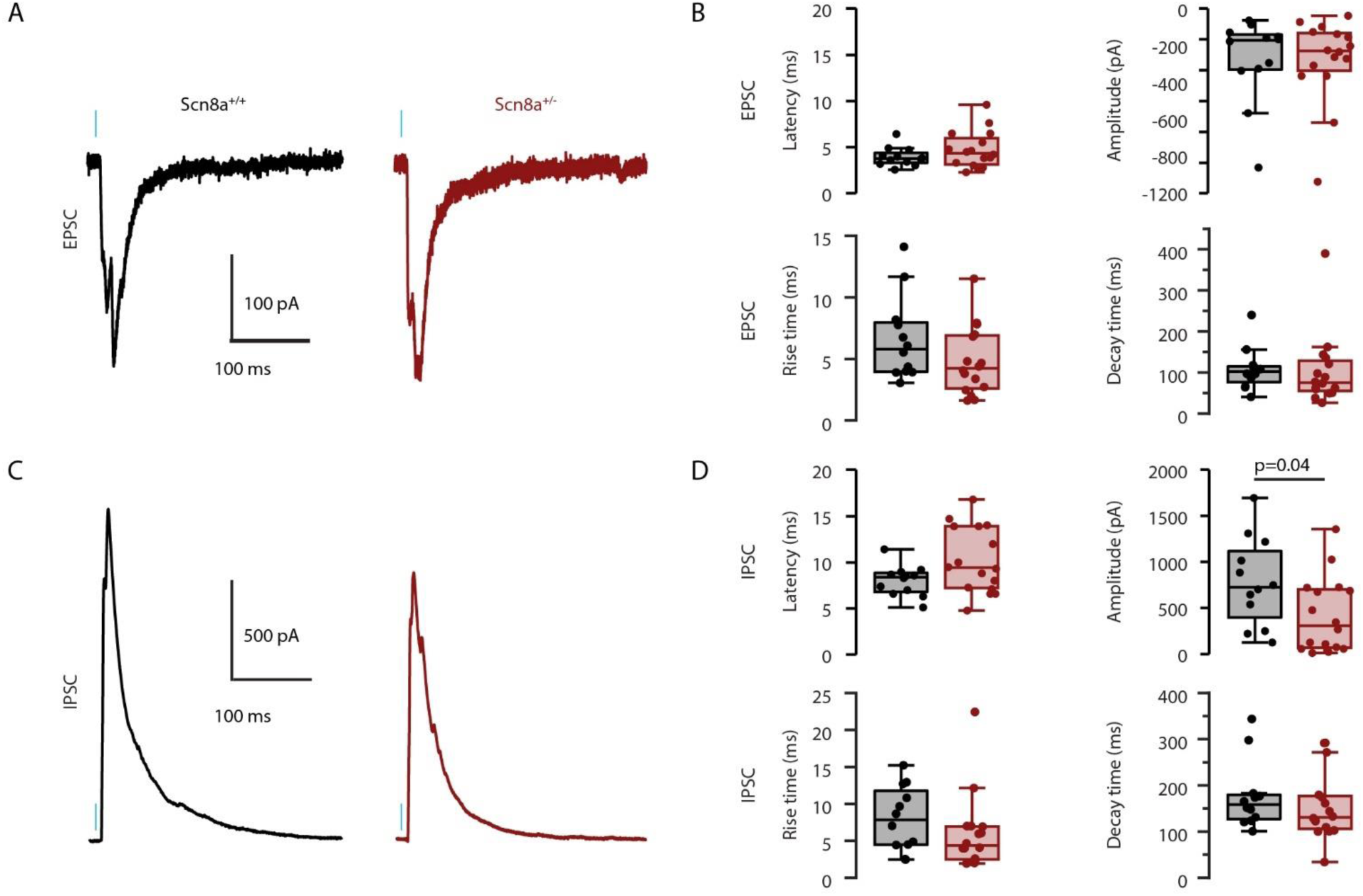
Synaptic responses in L5 pyramidal neurons evoked by Re afferent stimulation. (A) EPSCs recorded at -70 mV in L5 pyramidal neurons from mice expressing ChR2 in Re afferents. (B) Quantification of EPSC properties. (C) IPSCs recorded at +10 mV in L5 pyramidal neurons. (D) Quantification of IPSC properties.

**Suppl. Fig. 2.**
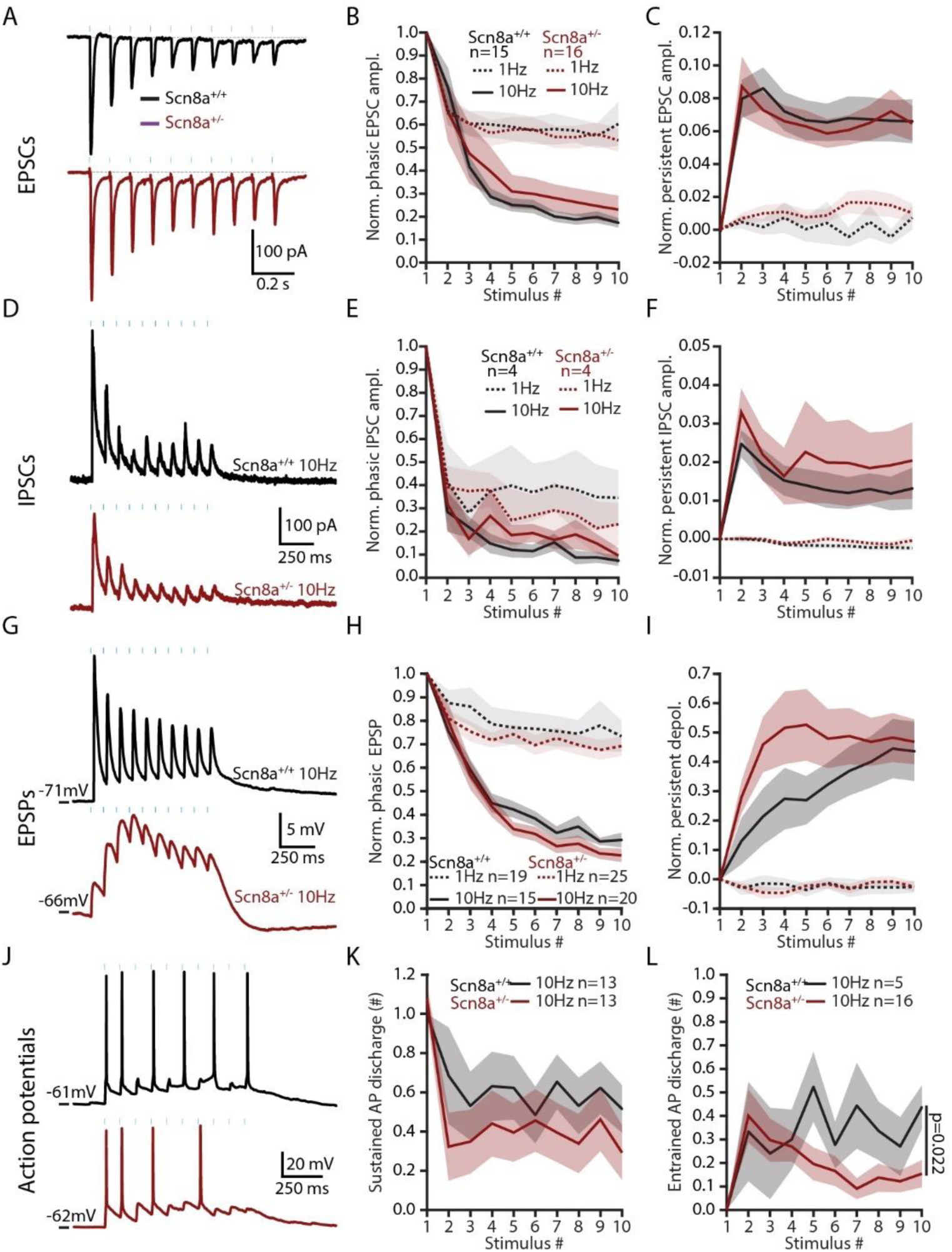
Synaptic and spiking responses of L5 pyramidal neurons to repeated Re afferent stimulation. (A) EPSCs evoked during 10 Hz train stimulation of Re afferents. (B) Quantification of phasic EPSC amplitude normalized to the first EPSC. (C) Quantification of persistent EPSC amplitude normalized to the first EPSC. (D–F) Same as (A–C) for IPSCs. (G–I) Same as (A–C) for subthreshold EPSPs. (J) Action potential discharge from resting membrane potential during 10 Hz train stimulation of Re afferents. (K) Quantification of sustained action potential firing. (L) Quantification of entrained action potential firing.

**Suppl. Fig. 3.**
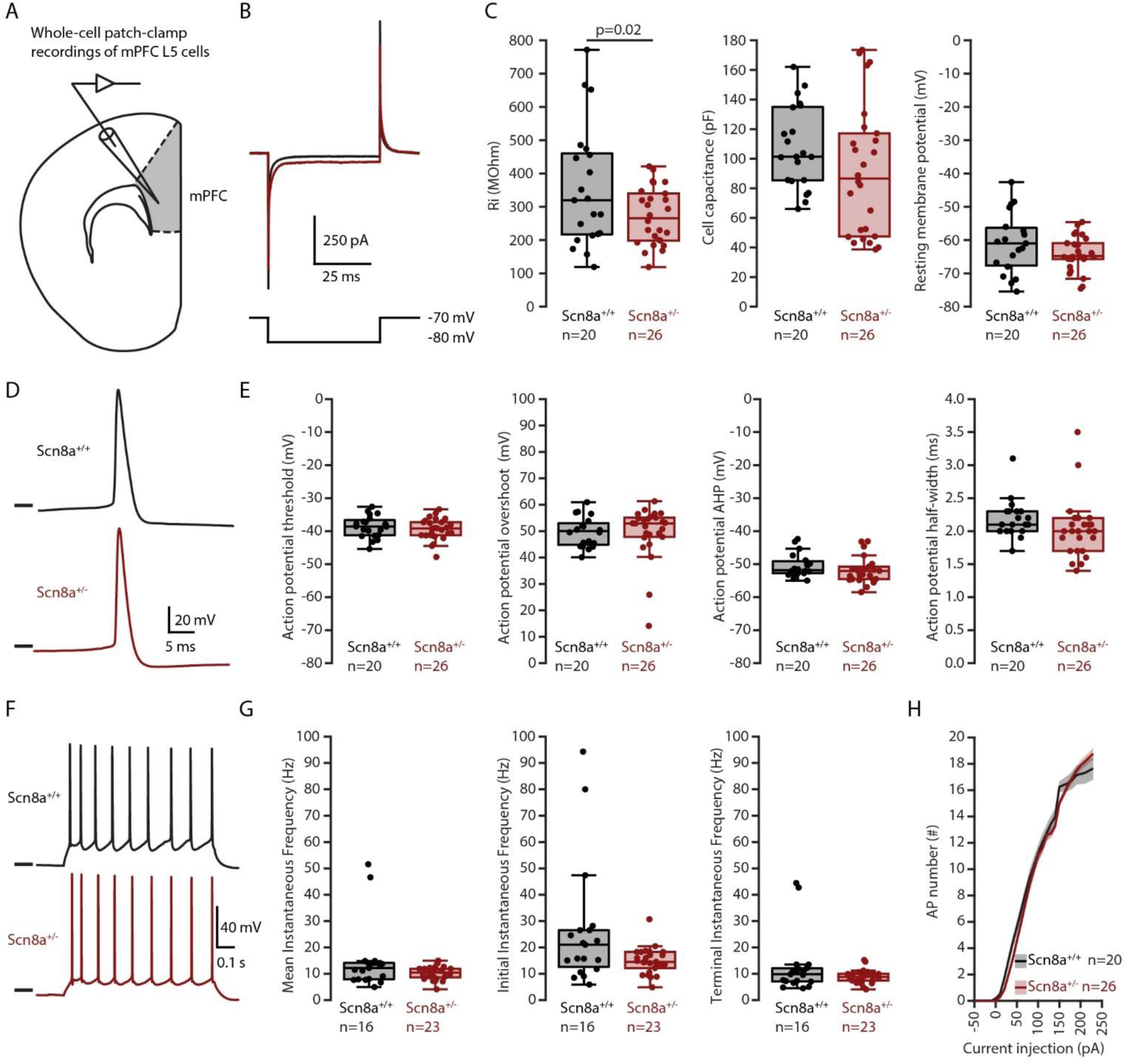
Cellular properties of mPFC L5 pyramidal neurons. (A) Whole-cell patch-clamp recordings from L5 pyramidal neurons in mPFC slices. (B) Membrane response to a 10-mV hyperpolarizing step. (C) Quantification of input resistance (Ri), cell capacitance, and resting membrane potential. (D) Representative action potentials. (E) Quantification of action potential threshold, overshoot, afterhyperpolarization, and half-width. (F) Action potential discharge during positive current injection. (G) Quantification of mean, initial, and terminal instantaneous firing frequency at twice the rheobase. (H) Quantification of the number of action potentials evoked during 1-s current injections of increasing intensity.

**Suppl. Fig. 4.**
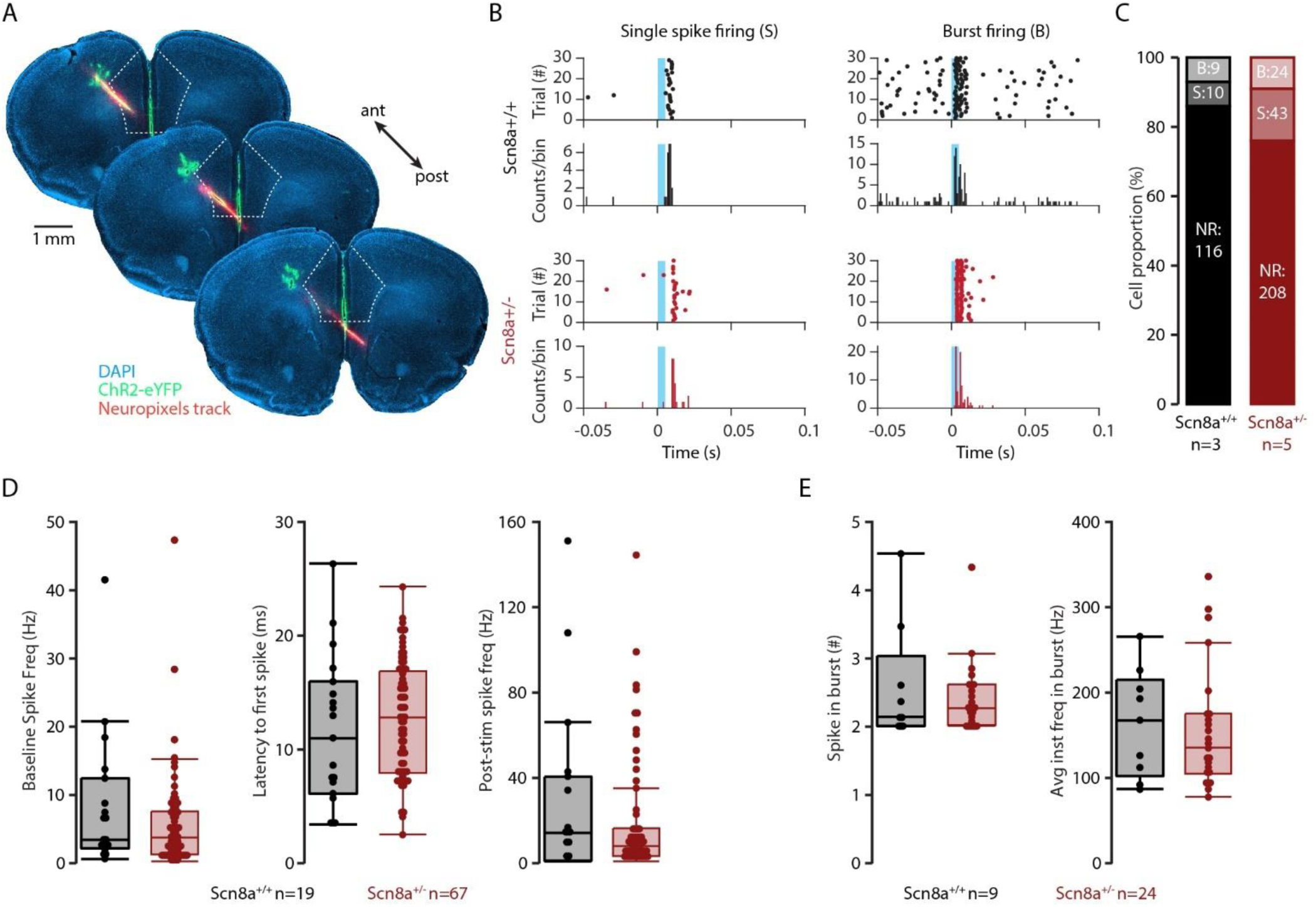
Re optogenetic stimulation drives spiking of mPFC units in vivo. (A) Epifluorescent images of mPFC coronal slices showing ChR2-eYFP–labeled Re afferents (green), Neuropixels probe tracks (red), and DAPI-stained nuclei (blue). (B) Raster plots and cumulative histograms from Scn8a^+/+^ (black) and Scn8a^+/-^ (red) mice showing single-spike firing (left) and burst firing (right) of mPFC units following Re stimulation. (C) Proportion of mPFC units that did not respond (NR) or responded with a burst (B) or single spike (S). (D) Quantification of baseline firing frequency, latency to first spike after laser onset, and post-stimulation firing frequency within 0–30 ms. (E) Quantification of the number of spikes per burst and the average instantaneous frequency within bursts.

**Suppl. Fig. 5.**
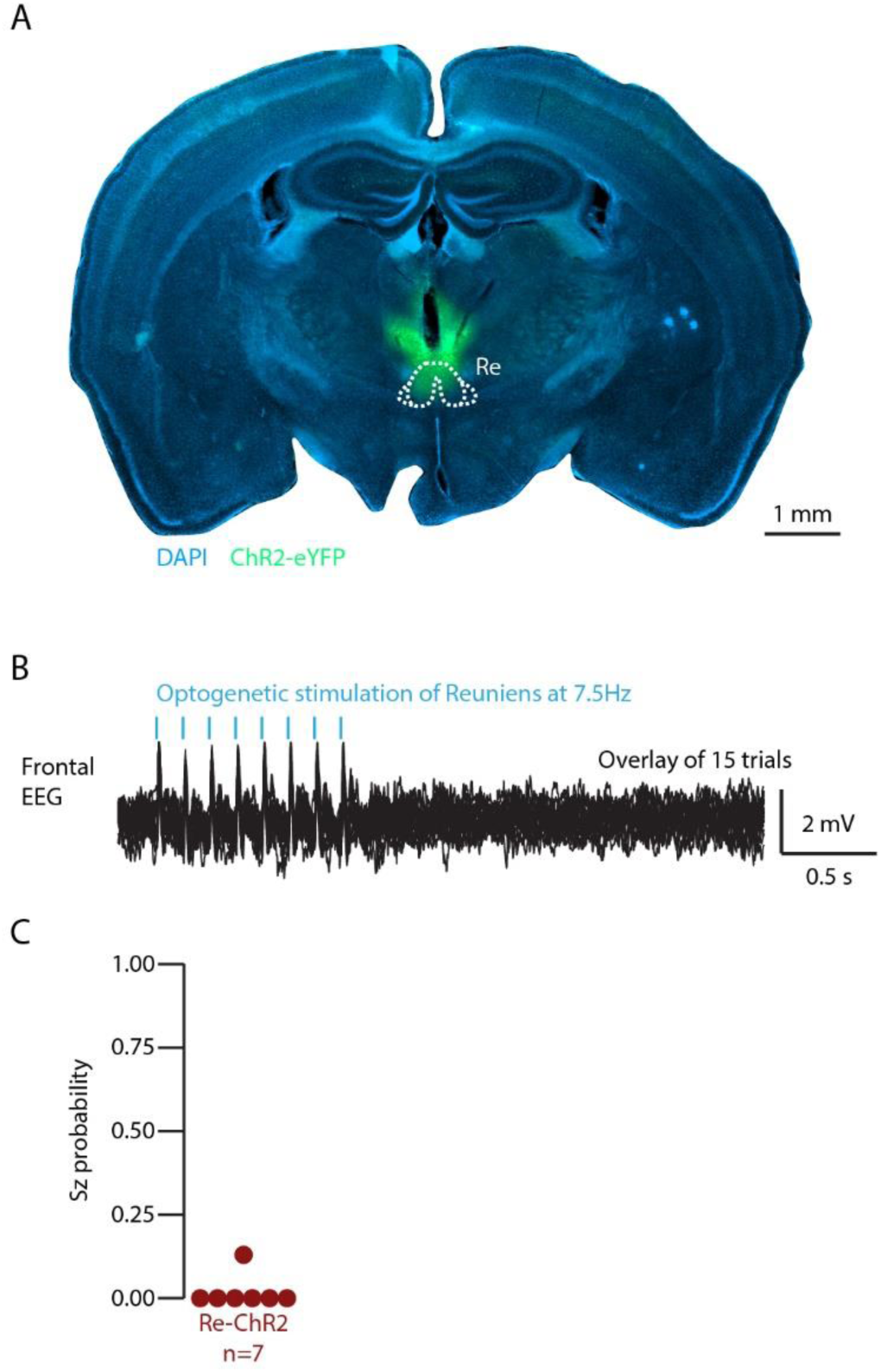
Stimulation of Re at 7.5 Hz, the main seizure frequency, does not drive SWDs. (A) Micrograph of a coronal brain slice with ChR2-eYFP (green) expression in Re, and DAPI-stained nuclei (blue). (B) Overlay of 15 traces showing brief deflection in the frontal EEG signal upon optogenetic stimulation of Re at 7.5 Hz. (C) Quantification of Seizure probability for each mouse during 7.5 Hz stimulation of Re.

